## Supplemental Figures and Tables for "*Ustilago maydis* Trf2 ensures genome stability by antagonizing Blm-mediated telomere recombination: fine-tuning DNA repair factor activity at telomeres through opposing regulations"

**Supplementary Figures 1S – 7S**

**Supplementary Tables 1S – 2S**

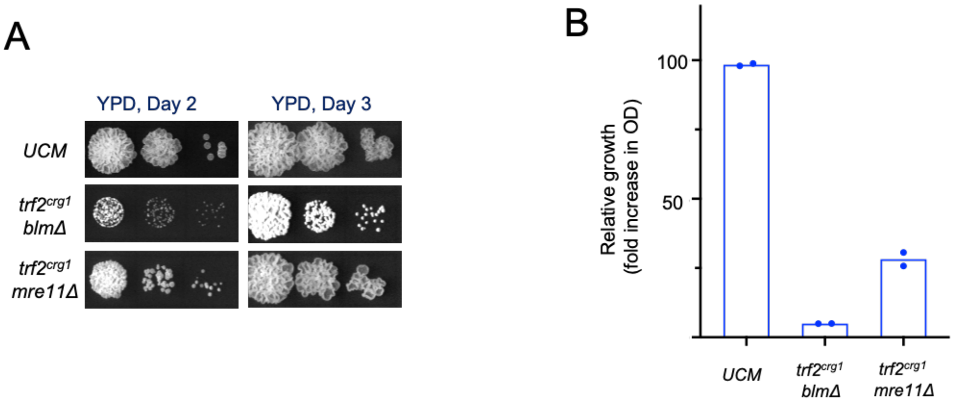

**Fig. 1S. Growth defects of the *trf2$^{crg1}$ blm*Δ and *trf2$^{crg1}$ mre11*Δ mutants grown in YPD**
**A.** Serial dilutions of the indicated strains were spotted onto YPD medium. The growth of the strains was imaged after 2 days and 3 days.
**B.** The indicated strains were inoculated into fresh YPD at an $OD_{600}$ of 0.01 and grown at 30 degree for 17 hours. The fold increases in $OD_{600}$ for the cultures were determined and plotted.

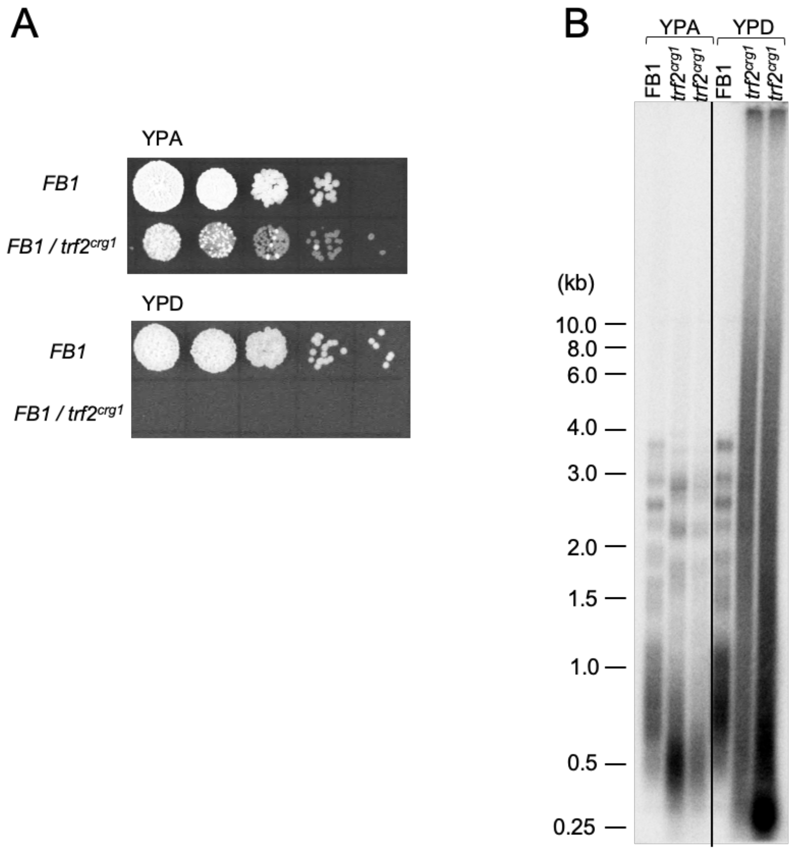

**Fig. 2S. Phenotypic analysis of Trf2-deficient cells derived from the FB1 strain background**
**A.** Serial dilutions of the indicated strains were spotted onto YPA and YPD media and incubated at 30ºC. Following 2 days of growth, the plates were imaged.
**B.** Genomic DNAs from the indicated strains were subjected to TRF Southern analysis.

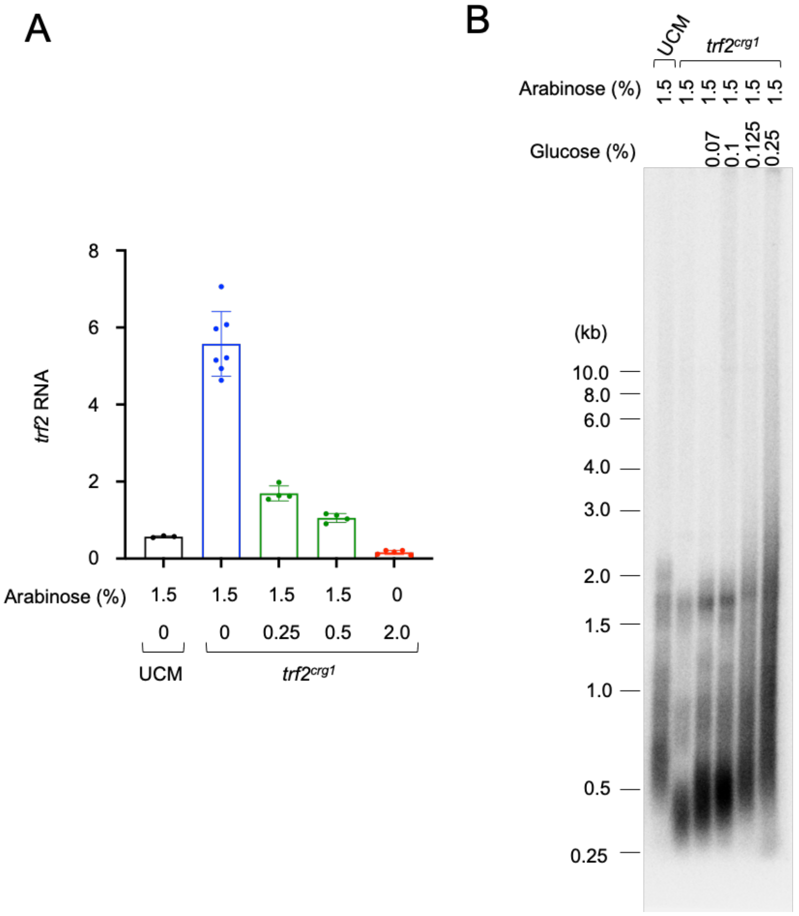

**Fig. 3S. A progressive reduction in Trf2 expression level correlates with a gradual increase in telomere lengths**

**A.** UCM and *trf2*$^{crg1}$ were grown in media containing 1.5% arabinose and varying concentrations of glucose (to reduce the activity of the *crg1* promoter). RNAs were isolated from these cultures and subjected to RT-qPCR analysis to determine the relative *trf2* RNA levels.

**B.** Genomic DNAs were isolated from UCM and *trf2*$^{crg1}$ grown in media with the specified arabinose and glucose concentrations and subjected to TRF Southern analysis.

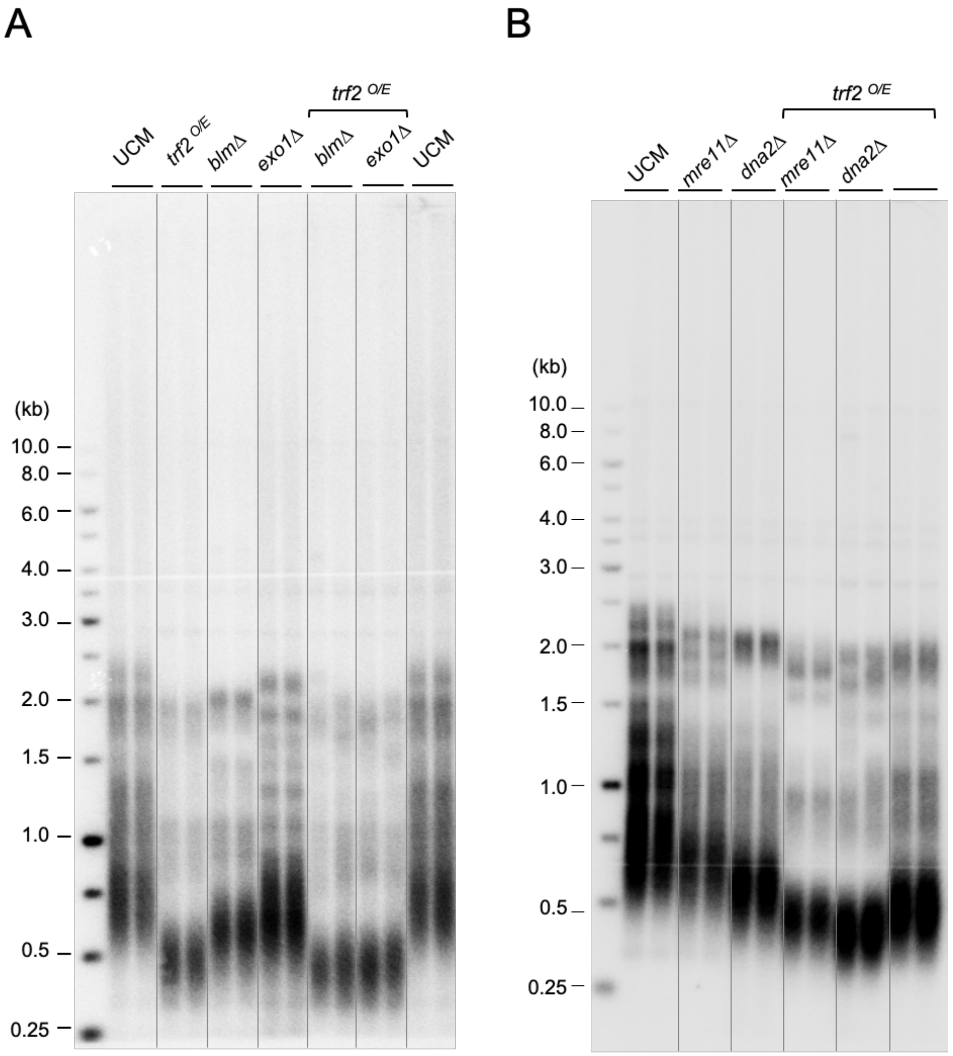

**Fig. 4S. Epistasis analysis of telomere shortening mediated by Trf2-overexpression and deletions of *exo1*, *blm*, *mre11*, and *dna2***

**A and B.** Genomic DNAs were isolated from the indicated strains grown in YPA, digested with *Pst*I, and subjected to TRF Southern analysis.

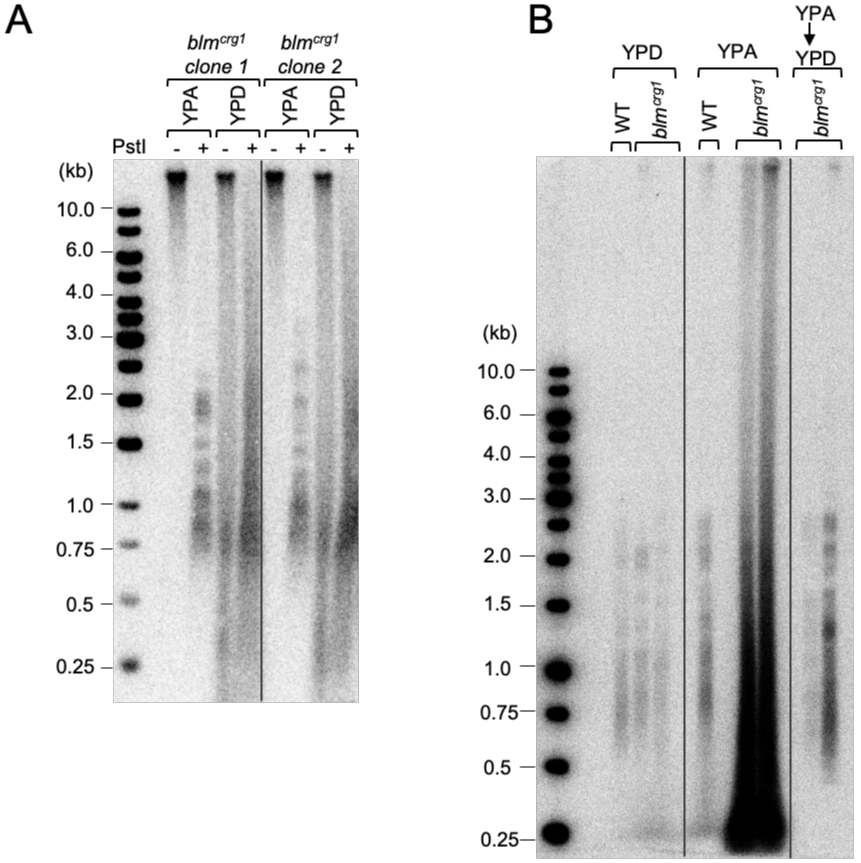

**Fig. 5S. Blm overexpression triggers ECTRs in a reversible manner.**
**A.** DNAs were isolated from *blm*$^{crg1}$ strains grown in the specified culture media and subjected to Southern analysis for telomere DNA with and without prior *Pst*I digestion.
**B.** DNAs were isolated from UCM and *blm*$^{crg1}$ strains grown in the designated culture media and subjected to Southern analysis. To test the reversibility of telomere defects induced by Blm overexpression, the *blm*$^{crg1}$ clones were first grown on a YPA plate and then re-streaked once on a YPD plate prior to liquid culture growth and genomic DNA isolation (designated by YPA -> YPD).

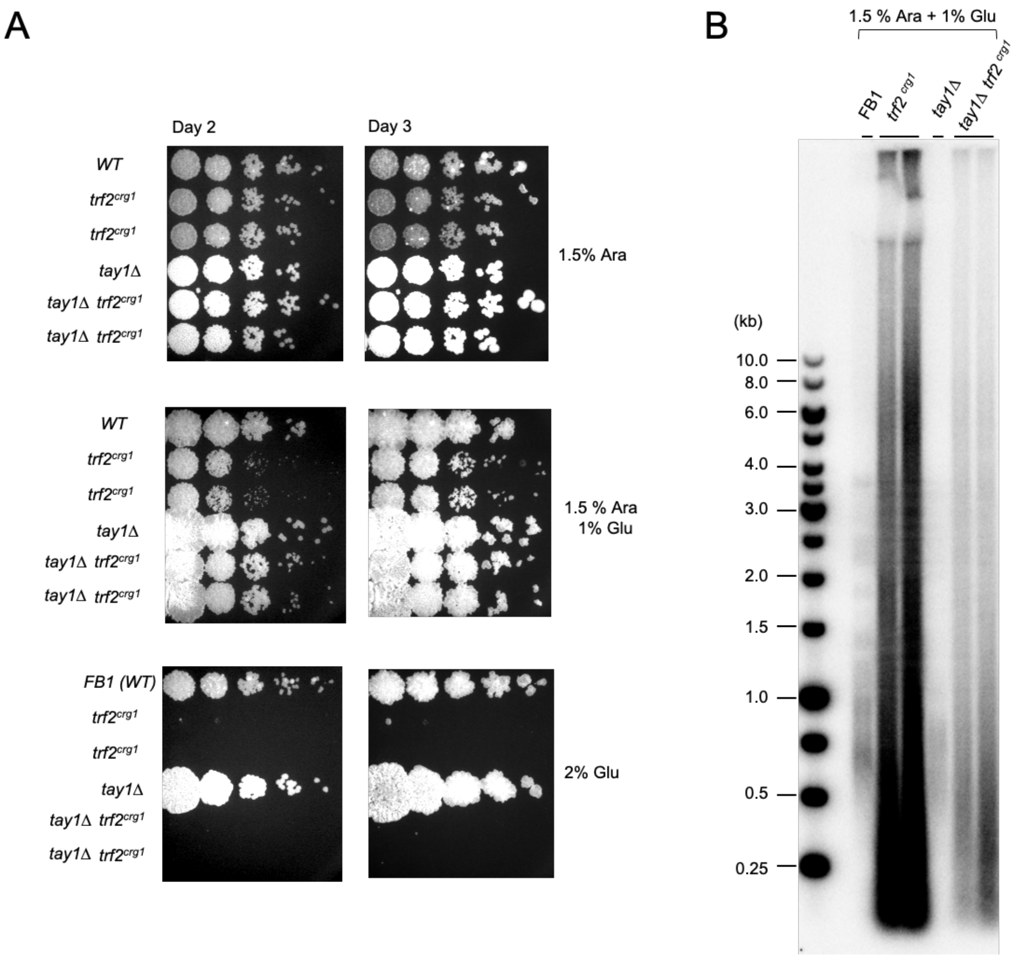

**Fig. 6S. Partial suppression of Trf2-deficiency phenotypes by *tay1Δ*.**

**A.** Serial dilutions of the designated strains were spotted onto semi-solid media containing the specified combinations of arabinose and glucose, and grown for 2 or 3 days at 30°C.

**B.** Genomic DNAs were isolated from the indicated strains grown in 1.5% arabinose and 1 % glucose, and subjected to TRF Southern analysis.

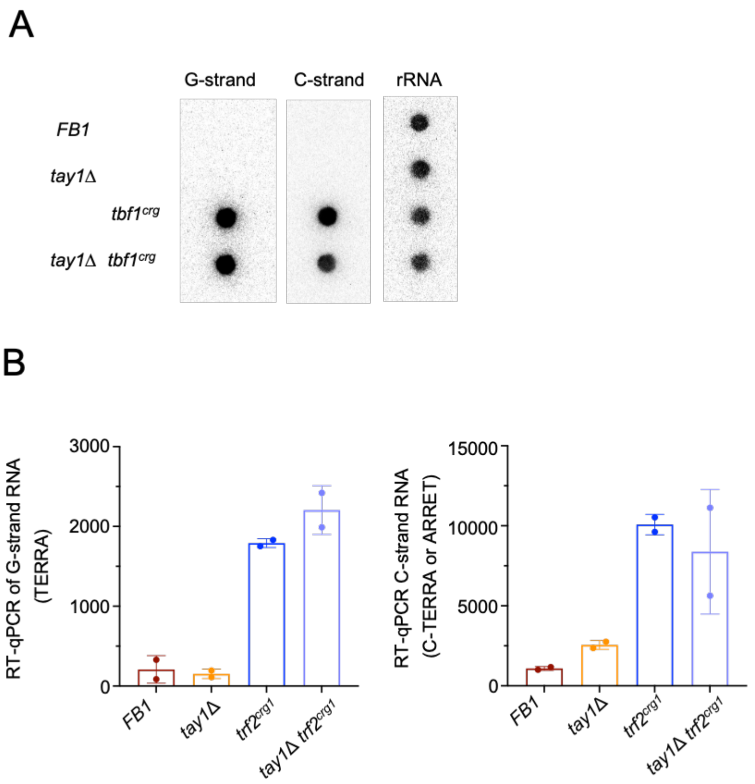

**Fig. 7S. Up-regulation of telomere G- and C-strand RNA synthesis in Trf2-deficient mutants**

**A.** RNAs were isolated from the indicated strains grown in YPD and spotted onto nylon membrane. After crosslinking, the membrane was probed sequentially for G-strand RNA, C-strand RNA, and rRNAs.

**B.** RNAs were isolated from the indicated strains grown in YPD, and subjected to RT-qPCR analysis to quantify the levels of G- and C-strand RNAs. For the G-strand RNA, the primers were designed to amplify TERRA (RNAs spanning subtelomeres and telomeres) from the UT6-bearing telomeres. For C-strand RNA, the primers can in principle amplify both C-TERRA (RNAs spanning subtelomeres and telomeres) and ARRET (RNAs with subtelomere sequences only) from UT6 telomeres. The quantities plotted represent cDNA copy numbers from 1 µl of RT reactions.

**Supplementary Table 1S. *U. maydis* strains used in this study**

| **Alias (Haploids)** | ***Relevant Genotype*** | **Reference** |
|---|---|---|
| FB1[a] | Wild type | (31) |
| USZ111 [ab] | *trf2* $^{crg1}$ | This work |
| UEY25 [ac] | *tay1*Δ | Yu et al., 2020 |
| USZ112 [abc] | *tay1*Δ *trf2* $^{crg1}$ | This work |
| UCM350 [d] | wild type | Kojic et al., 2002 |
| USZ100 [bd] | *trf2* $^{crg1}$ | Yu et al., 2020 |
| USZ121 [bde] | *blm*Δ *trf2*$^{crg1}$ | This work |
| USZ122 [bdf] | *dna2*Δ *trf2*$^{crg1}$ | This work |
| USZ123 [bdg] | *exo1*Δ *trf2*$^{crg1}$ | This work |
| USZ124 [bdh] | *mre11*Δ *trf2*$^{crg1}$ | This work |
| USZ125 [bdi] | *trf2*$^{crg1}$ *rad51*Δ | This work |
| USZ131 [j] | *blm*$^{crg1}$ | This work |
| USZ132 [j] | *blm*$^{crg1}$ | This work |
| USZ141 [k] | *blm*$^{nar1}$ | This work |
| USZ142 [bk] | *trf2*$^{crg1}$ *blm*$^{nar1}$ | This work |
| USZ143 [kl] | *pot1*$^{crg1}$ *blm*$^{nar1}$ | This work |

[a] The genotype of FB1 is *a1b1* for the mating type loci.
[b] *trf2* was placed downstream of the arabinose-dependent *crg1* promoter through the introduction of a Cbx$^R$-containing cassette.
[c] *tay1* was disrupted by the insertion of *hph* cassette expressing the hygromycin resistance gene *(Hyg$^R$).*
[d] The genotype of UCM350 *is nar1-6 pan1-1 a1b1*. *nar*, *pan*, and *ab* indicate inability to reduce nitrate, auxotrophic requirement for pantothenate, and mating type loci, respectively.
[e] *blm* was disrupted by the insertion of *hph* cassette expressing the hygromycin resistance gene *(Hyg$^R$).*
[f] *dna2* was disrupted by the insertion of *hph* cassette expressing the hygromycin resistance gene *(Hyg$^R$).*
[g] *exo1* was disrupted by the insertion of *hph* cassette expressing the hygromycin resistance gene *(Hyg$^R$).*
[h] *mre11* was disrupted by the insertion of *hph* cassette expressing the hygromycin resistance gene *(Hyg$^R$).*
[i] *rad51* was disrupted by the insertion of *hph* cassette expressing the hygromycin resistance gene *(Hyg$^R$).*
[j] *blm* was placed downstream of the arabinose-dependent *crg1* promoter through the introduction of a Cbx$^R$-containing cassette.
[k] *blm* was placed downstream of the nitrate-dependent *nar1* promoter through the introduction of a Hyg$^R$-containing cassette.
[l] *pot1* was placed downstream of the arabinose-dependent *crg1* promoter through the introduction of a Cbx$^R$-containing cassette.

**Supplementary Table 2S. Oligos used in this study**

| Name | Sequence 5' to 3' |
|---|---|
| **Protein Expression** | |
| UmTrf2-416F-SalI | ATTAC GTCGAC CG CAA TCC GAA CAG CGG TTA |
| UmTrf2-dn-st-NotI | AA GCGGCCGC CTA TTC GTT TGA AAG AGA ACT CGA CG |
| UmBlm-F-Nco | AAT CCATGG CA ATG CCG CAA TCC GCA CTA ACC CCA |
| UmBlm-R-FG-NotI | AAT GCGGCCGC CTA CTT GTC ATC GTC ATC CTT GTA ATC ACC CGA ACG AGG TAG ATT GGG C |
| **Strain Construction and Genotyping** | |
| UmBlm(1)Nde | GAAATC CAT ATG CCG CAA TCC GCA CTA |
| UmBlm(720R)Xba | ATT TCTAGA CAA GAT CTG AAA CGA GCT CG |
| UmBlm(-720)Xba | ATA TCTAGA TTG GAA CTG CGC AGT AAG CT |
| UmBlm(-1R)Eco | ATT GAATTC AAC TGA AAG AAG GCA ATC CT |
| UmBlm(720R)Nsi | AGA ATGCAT CAA GAT CTG AAA CGA GCT CG |
| UmBlm(-720)Nsi | ATA ATGCAT TTG GAA CTG CGC AGT AAG CT |
| **Helicase assays** | |
| NT – top | TTCTTCCTTTCCCTCTTCCTGATACGGCTGCTTCTCATCTACAACGTGATCCGTCATGGT |
| NT – bottom | ATGAGAAGCAGCCGTATCAGGAAGAGGGAAAGGAAGAA |
| Telo – top | TTCTTCCTTTCCCTC**TAGGGTTAGGGTTAGGGTTAGGGTTAGGGTTAGGGTTAGGGTTAG** |
| Telo – bottom | **CCCTAACCCTAACCCTAACCCTA**GAGGGAAAGGAAGAA |
| **PCR, hybridization, and EMSA assays** | |
| $TTAGGG_4$ (G4) | TTAGGG TTAGGG TTAGGG TTAGGG |
| $CCCTAA_4$ (C4) | CCCTAA CCCTAA CCCTAA CCCTAA |
| $TTAGGG_8$ (G8) | TTAGGG TTAGGG TTAGGG TTAGGG TTAGGG TTAGGG TTAGGG TTAGGG |
| $CCCTAA_8$ (C8) | CCCTAA CCCTAA CCCTAA CCCTAA CCCTAA CCCTAA CCCTAA CCCTAA |
| UmrRNA_26S_121F | GCTTCGGACCATGCCTAAG |
| UmrRNA_26S_642R | CTTGGTCCGTGTTTCAAGACG |
| **RT-PCR and RT-qPCR** | |
| UT6-TERRA-F1 | GGACGGCAGATATATATTGTGAGTGG |
| UT6-TERRA-F2 | GTGGCAACATTGGGTGAGC |

| | |
|---|---|
| UT6-TERRA-R1 | CCGTTGACACATTCAATCCCTC |
| UT6-TERRA-R2 | CTTCAAGCCCTGCAGCC |
| CCCTAA$_4$ (C4) | CCCTAA CCCTAA CCCTAA CCCTAA |
| Blm-PCR-180F | CAC TGC TCA AAA GAG ATC GAG GTC TGG |
| blm-PCR-209F | CAT CGG CCG CTT CCA TCT CAA ACG |
| blm-PCR-305R | CGA GGT GTC TGC GAT ATC GAG CCA C |
| Blm-PCR-400R | CAA AAT CGA TCC GAA GCT CGT CTT CG |
| Blm-PCR-500R | AAG GCT TCG TGC TTT TGG TAC CAA G |
| Trf2-PCR-1100R | TCT TCC GAG CCT TCA TCC GA |
| Trf2-PCR-865-F | CGC GAC ACC AAC CAT ACA AGC A |
| Trf2-PCR-939-R | GGT GAG TGG GGC ACT GTG TC |
| **STELA and fusion assays** | |
| UT4-F | TCGGGCAACGTTCCATGTCG |
| UT4-subtel-R2375 | CCCTCGAAGGCAGTGCATAC |
| UT6-F | CTACTACACATCGGTTCAGGC |
| UT6-subtel-R2400 | ATGCCAAAGTGGAAATCGTGCAC |
| C Telorette 1 | GCTCCGTGCATCTGGCATCCCCTAAC |
| C Telorette 2 | GCTCCGTGCATCTGGCATCTAACCCT |
| C Telorette 3 | GCTCCGTGCATCTGGCATCCCTAACC |
| C Telorette 4 | GCTCCGTGCATCTGGCATCCTAACCC |
| C Telorette 5 | GCTCCGTGCATCTGGCATCAACCCTA |
| C Telorette 6 | GCTCCGTGCATCTGGCATCACCCTAA |
| Teltail | GCTCCGTGCATCTGGCATC |
